# Combining chemogenomic and gene-dose assays to investigate drug synergy

**DOI:** 10.1101/2021.02.01.429253

**Authors:** Hamid Gaikani, Andrew M. Smith, Anna Y. Lee, Guri Giaever, Corey Nislow

**Affiliations:** Department of Pharmaceutical Sciences, University of British Columbia, Vancouver BC V6T 1Z3, Canada; Department of Chemistry, University of British Columbia, Vancouver BC V6T 1Z1, Canada; Department of Biochemistry and Molecular Biology, University of British Columbia, Vancouver BC V6T 1Z1, Canada; University of Toronto, Donnelly CCBR, Toronto, ONT, M5S 3E1, Canada

**Keywords:** drug synergy, drug combinations, drug-gene interaction, antifungal

## Abstract

From the earliest days of using natural remedies to modern applications of clinically tested medications, combining therapies for disease treatment has been standard practice. Combination treatments can exhibit synergistic effects, broadly defined as a greater-than-additive effect of two or more therapeutic agents. Indeed, clinicians often use their experience and expertise to tailor such combinations in the hopes of maximizing the therapeutic effect. Alongside these efforts, computational studies into understanding and predicting the biophysical underpinnings of how synergy is achieved have benefitted from high-throughput screening and computational biology. One challenge is how to best design and analyze the results of synergy studies performed at scale, especially because the number of possible combinations to test quickly becomes unmanageable, and the tools to analyze the resulting data are quite new. Nevertheless, the benefits of such studies are clear — by combining multiple drugs in the treatment of infectious disease and cancer, for instance, one can lessen host toxicity and simultaneously reduce the likelihood of resistance to treatment. In this study, we extend the widely validated chemogenomic HIPHOP assay to drug combinations. We identify a class of “combination-specific sensitive strains” that suggest mechanisms for the synergies we observe and further suggest focused follow-up studies.

## Introduction

Drugs and drug-like molecules are powerful molecular tools that can act by rapid and reversible inhibition of a specific protein or other biomolecule in cells. Such chemical perturbations, while similar to genetic manipulations, have several experimental advantages: they are tunable, fast-acting, often reversible, and can be applied across large evolutionary distances, e.g., from yeast to human. Drugs can be easily combined to simultaneously modulate multiple proteins’ activities, and in fact, the modulation of gene products by administering a combination of drugs can be vital for a successful course of treatment^1^. The clinical success of chemical combination therapies has motivated our empirical study of synergistic chemical interactions. These data can then be assessed to predict how two drugs might interact in a biological system. To study the potential interaction, several mathematical models of drug synergy are available^2–4^; two widely used approaches are the Bliss model of independence^3^ and the Loewe additivity model^4,5^. Neither model is able to explain all drug synergies, and no mathematical model is suited for all observed chemical interactions, indicating the complexity of the problem.

Invasive fungal infections (IFI) are lethal threats to human health, and they cause almost two million worldwide deaths annually. In 2018, the death rate among patients suffering from IFI was reported to be 28.8%^6^. At present, the available therapies, particularly for invasive infections, are limited to four categories of antifungal drugs; azoles, polyenes, echinocandins, and 5-flucytosine^7^, and the clinical results from most IFI cases are not optimal. In addition, emerging pathogens resistant to common antifungals^8^ such as the pan drug-resistant yeast Candida auris have spread in health care facilities globally^9,10^. One potential solution to the dearth of effective treatments is to explore the antifungal efficacy of novel drug combinations, including those prescribed for diverse indications^11^. The use of drug combinations gives rise to several opportunities: 1) it has been proven, both empirically and theoretically, that drugs that are synergistic for a particular effect do not tend to show synergy for side effects^12^ 2) the dose of individual agents with serious side effects can be reduced in a combination 3) synergistic antifungal activity increases therapy potency and reduces lengths of treatment and 4) compared with monotherapy, it minimizes the risk for antifungal resistance^11^.

Considering the limited number of drugs available for IFI treatment^13^, we sought to expand upon our strategy to use yeast as a eukaryotic model to screen any drug that inhibits the growth of, or kills yeast. Even though such drugs may be active against the host itself, our rationale is that using these drugs could lower host-dependent side-effects because each agent in a combination is typically applied in lower doses. In this study, we selected 10 compounds based upon their well-characterized targets in yeast, and from 100 possible combinations, drug pairs that empirically showed synergy were used in HIP–HOP assays — a validated genome-wide screen based on Haploinsufficiency Profiling (HIP) and Homozygous Profiling (HOP) to quantify the relative abundance of uniquely tagged yeast deletion strains. We found that, using this approach, we were able to; 1) identify numerous synergistic combinations, 2) quantify this synergy and identify combination-specific sensitive strains on a genome-wide scale.

## Results

### Synergy screens

Using our database of drugs and drug-like molecules^14^ we selected 10 compounds (Table 1) and screened all possible combinations of these drugs in a 6-by-6 dosage matrix using growth curve analysis (Figure 1). The growth data were analyzed, using a computer program (AUDIT)^15^ that converts raw absorbance values into growth curves. These curves were then analyzed to generate a heatmap^16^, examples of which are shown in Figures 1 and 2. Next, to determine potential synergy, each heat map and corresponding growth curve data were examined to produce the average epsilon score for the drug combination 6-by-6 matrix (defined as AvgS). If two individual drugs act independently, their effects are expected to combine multiplicatively. In other words, if a drug affecting gene x causes a fitness effect W_x_, and a drug affecting gene y causes a fitness effect W_y_, then the total effect of the drug combination (W_xy_) is predicted to be W_x_ × W_y_. For our purposes, we measured the deviation epsilon (ε) from this expectation (where εxy = W_xy_ – W_x_ × W_y_)^17,18^. Using this score and a threshold of AvgS < −0.05 to score a combination as synergistic, 33% of all combinations showed synergy. Given this unexpectedly high level of synergy, we applied additional filter; namely that three other drug combination models (additivity^4,5,19^, highest single agent, and a potentiation model^2^) identified a synergistic combination, we identified 10 combinations that deviated from expectation in all 4 models and which were therefore classified as synergistic. Interestingly, fenpropimorph vs. miconazole, fenpropimorph vs. cerivastatin, and miconazole vs. cerivastatin all possessed a strong synergy when combined. These are the only compound pairs that target the same pathway — consistent with the idea that drugs targeting the same essential pathway can be effective means to produce synergistic combinations.

**Table 1.** Compounds used and known targets

| Compound | Protein Target | Method |
| --- | --- | --- |
| cerivastatin | Hmg1<sup>20</sup> | Bar-seq |
| tunicamycin | Alg7<sup>21</sup> | MSP/HIP |
| methotrexate | Dfr1<sup>22</sup> | HIP |
| miconazole | Erg11<sup>23,24</sup> | HIP |
| rapamycin | Tor2<sup>25,26</sup> | MSP/HIP |
| cantharidin | Glc7<sup>27,28</sup> | HIP |
| fenpropimorph | Erg24<sup>29,30</sup> | HIP |
| latrunculin A | Act1<sup>31,32</sup> | Resistance mapping |
| benomyl | Tub1<sup>33,34</sup> | HIP and mutant mapping |
| sodium fluoride | Ipp1<sup>35</sup> | HIP |
| hydroxyurea | Rnr1<sup>36,37</sup> | HIP and resistance mapping |

**Figure 1.**
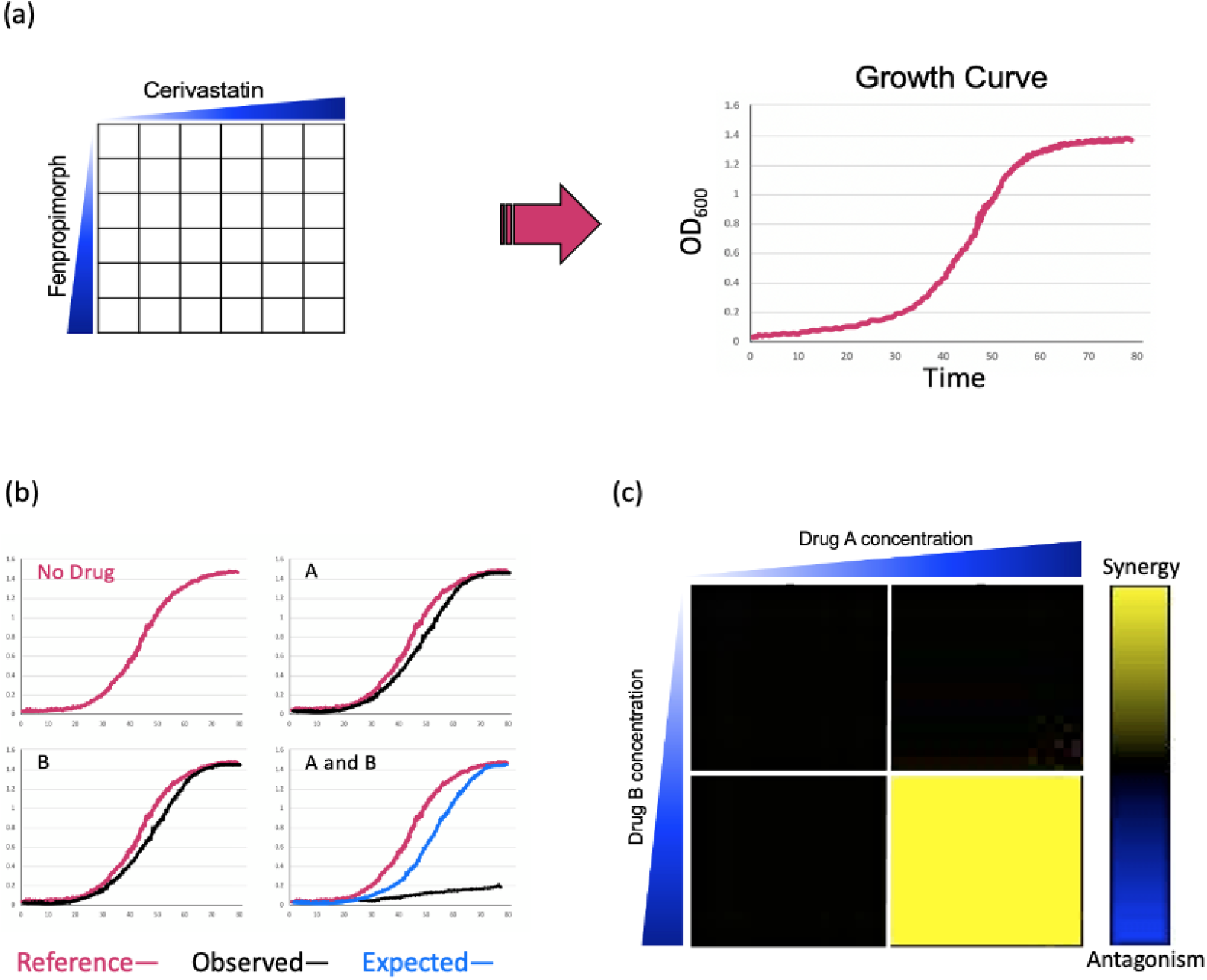
Combination screening and transformation of growth data into a quantitative metric. (a) illustrates the combination of fenpropimorph vs cerivastatin as an example. A 6-by-6 dose matrix is screened with an increase of each drug along the x and y axes. The blue triangles represent increasing concentration of drug. Each square in the 6-by-6 matrix is represented by a growth curve for each drug combination, which is optical density (OD_600_) vs. time. (b) To quantify synergy, growth data (a) were transformed into heat map (b). The red growth curve is the DMSO (no drug) control. The cells marked A and B, represent the addition of compound A and compound B, respectively. The black line represents the growth in each drug condition. The combination of A and B is shown in the bottom right-hand cell in which the blue line indicates the expected growth rate based on the multiplicative model, while the black curve is the actual cell growth in the drug combination. In the heatmap (c), the color of the square represents epsilon generated using the multiplicative model, where black represents no interaction, yellow represents synergy and blue represents antagonism.

**Figure 2.**
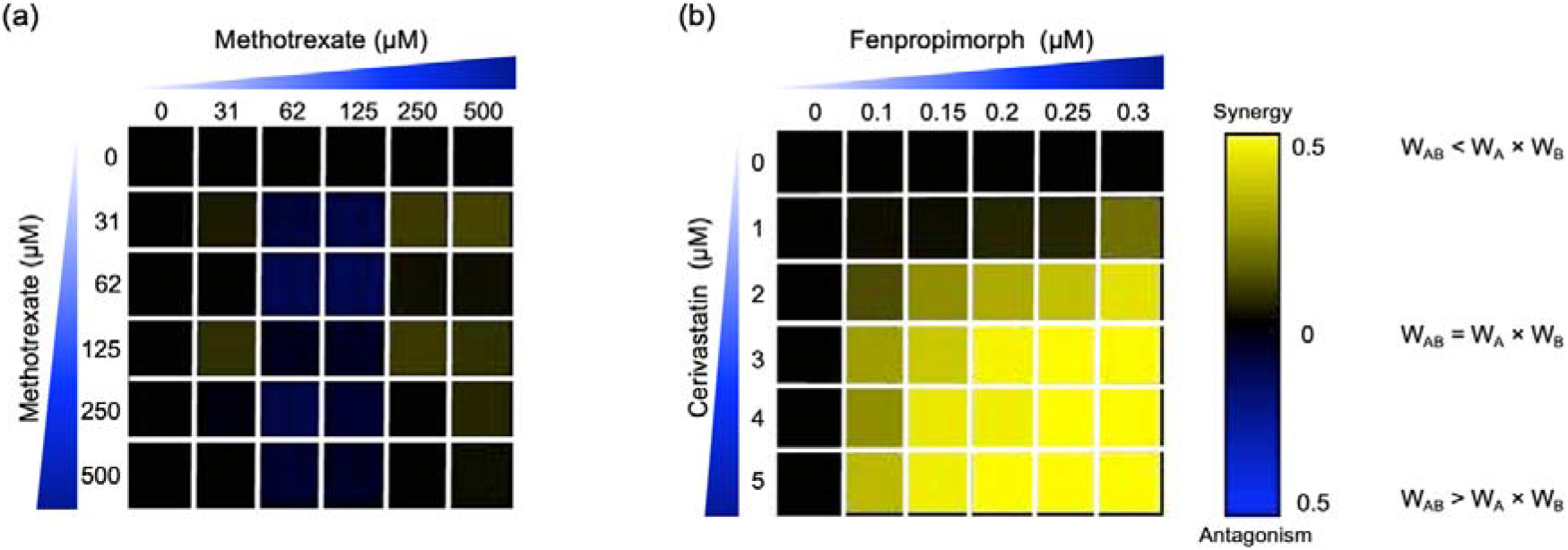
Two examples of drug combinations screened. The concentration of drug increases along a particular axis, and the color of the square represents epsilon generated by using the multiplicative model. (a) methotrexate vs. methotrexate is an example of independence. In heat map (a), we expect epsilon to be near zero as methotrexate vs. methotrexate should not be synergistic since the same compound is added on both axes. (b) In contrast the 6-by-6 dose response matrix of cerivastatin vs. fenpropimorph provides an example of synergy, because epsilon is negative. W_AB_ represents the relative growth of yeast in the presence of both drugs A and B when compared to a no-drug control, W_A_ represents the relative growth of yeast in the presence of drug A when compared to a no drug control. The color-coded scale bar from yellow (synergy) to blue (antagonism) covers the spectrum where W_ab_ < W_a_ × W_b_, to no interaction W_ab_ = W_a_ × W_b_ to antagonism W_ab_ > W_a_ × W_b_.

### Using synergistic drug combinations to predict drug-gene interactions

Having identified a high-confidence set of synergistic drug-drug interactions, we tested if these interactions could be recapitulated by combining the relevant drug-gene interactions. Our rationale was that if two drugs were synergistic, a loss-of-function mutation (as exemplified in the heterozygous state) in one of the known drug-targets would confer hypersensitivity to the second compound. In other words, if Drug A targets protein A and Drug B targets protein B to produce synergy, then one would predict that Drug A when combined with a loss-of-function mutant in B, should phenocopy the drug combination. To empirically test this prediction, we selected heterozygous deletion mutants of the known drug targets and challenged them with each drug listed in (Table 1), and the results were compared to data derived from the combination screen presented in Figure 1.

11 heterozygous deletion mutants — each deleted for one of the known drug-targets — along with a wild type control strain were profiled in each drug, results in 121 drug-gene interaction tests (i.e., 11 drugs against 11 heterozygous deletion mutants). The act1 heterozygote displayed a significant fitness defect without drug treatment and was eliminated from further analysis. The remaining 110 drug-gene interactions were examined for drug sensitivity. We used a cut-off of greater than a 10% fitness defect (i.e., an inhibitory concentration of 10 or IC10) and identified 72 negative drug-gene interactions. Among these 72 interactions, 10 were the expected HIP-drug interaction, while 62 negative drug-gene interactions were novel. In the 62 negative drug-gene interactions, 17 of the 18 predicted interactions had greater than a 10% defect in growth, giving a significant enrichment (P-value = 0.003) of drug-gene interactions, which are predicted by synergistic combinations. Using a more stringent cut-off of greater than a 30% fitness defect, 15 negative drug-gene interactions, 8 of which were from the predicted drug-gene interactions (P-value = 0.0002); Table 2.

**Table 2.** Fitness of heterozygous mutants in drugs. Red denotes expected drug-induced haploinsufficiency (e.g., when considering an ERG11 Het, the expectation is that this strain would grow slower than its wild-type counterpart in miconazole because miconazole targets the ERG11 protein.); yellow indicates predicted haploinsufficiency interactions based on synergistic drug combinations (i.e., based on the drug-drug synergy we expect a drug-gene interaction). Numbers represent the average generation time (AvgG) of the mutant’s fitness relative to wild type in the drug condition.

|  | miconazole<br>200nM | benomyl<br>0.02μg/mL | rapamycin<br>6nM | fenpropimorph<br>0.04% | cantharidin<br>250μM | fluoride 40mM<br>40 mM | hydroxyurea<br>80mM | methotrexate<br>450μM | cerivastatin<br>2μM | tunicamycin<br>0.2μM | latrunculin A<br>0.5μM |
| --- | --- | --- | --- | --- | --- | --- | --- | --- | --- | --- | --- |
| **erg11** | 0.82 | 0.76 | 0.94 | 0.54 | 0.95 | 1.00 | 0.92 | 0.99 | 0.90 | 0.99 | 0.90 |
| **tub1** | 0.72 | 0.81 | 0.90 | 0.42 | 0.69 | 0.87 | 0.78 | 0.83 | 0.81 | 0.82 | 0.35 |
| **tor2** | 0.53 | 0.73 | 0.56 | 0.68 | 0.90 | 0.98 | 0.85 | 0.96 | 0.87 | 0.89 | 0.65 |
| **erg24** | 0.55 | 0.74 | 0.71 | 0.73 | 0.88 | 0.89 | 0.83 | 0.90 | 0.77 | 0.87 | 0.68 |
| **glc7** | 0.52 | 0.79 | 0.65 | 0.50 | 0.58 | 0.95 | 0.83 | 0.90 | 0.88 | 0.83 | 0.68 |
| **ipp1** | 0.61 | 0.74 | 0.72 | 0.78 | 0.91 | 0.71 | 0.89 | 0.97 | 0.99 | 0.89 | 0.68 |
| **rnr1** | 0.64 | 0.92 | 0.73 | 0.56 | 0.93 | 0.96 | 0.87 | 0.94 | 0.90 | 0.93 | 0.68 |
| **dfr1** | 0.55 | 0.92 | 0.80 | 0.68 | 0.93 | 1.04 | 0.87 | 0.78 | 0.97 | 0.91 | 0.75 |
| **hmg1** | 0.56 | 1.07 | 0.80 | 0.68 | 0.97 | 1.05 | 0.90 | 1.01 | 0.75 | 0.91 | 0.87 |
| **alg7** | 0.76 | 1.05 | 0.81 | 0.55 | 0.99 | 1.05 | 0.94 | 1.04 | 0.96 | 0.79 | 0.96 |

### Determination of a background synergy rate

To put our drug-drug synergy observations in context, we sought to determine the chances of observing synergy when two randomly selected compounds were combined. In other words, establishing the likelihood observing synergistic effects of any two compounds would then allow one to calculate any enrichment over random chance. Accordingly, we tested all pairwise combinations of 15 compounds in a 4-by-4 dosage matrix (Table 3). Our criteria for selecting such “random compounds” included i) they were bioactive in yeast, and ii) their HIP–HOP profiles showed a similar number of sensitive strains when compared to the compounds in table 2. We further evaluated these compounds by mapping them onto the synthetic genetic array (SGA) network of gene-gene interactions^38^. Using the multiplicative synergy model, 17% of these ‘random’ combinations were synergistic (ε < −0.20) which dropped to 9.5% of combinations when the over-represented compounds that affect the cell wall and secretion were excluded. This value is similar to previously reported combination screening studies, which report ~10% baseline synergy in any combination^2,39,40^.

**Table 3.** Drugs and chemical probes and their concentrations used in a 4-by-4 dose matrix to determine baseline synergy. The three compounds indicated by their CAS numbers are from reference^14^.

| Drug | Yeast Target | Dose 1 | Dose 2 | Dose 3 | Dose 4 | Units |
| --- | --- | --- | --- | --- | --- | --- |
| cisplatin | DNA | 0 | 100.0 | 150.0 | 200.0 | μM |
| MMS | DNA | 0 | 0.136 | 0.318 | 0.454 | mM |
| camptothecin | TOP2 | 0 | 0.25 | 0.5 | 1.0 | mM |
| caspofungin | FKS1 | 0 | 7.0 | 7.2 | 7.5 | nM |
| chlorpromazine | unknown | 0 | 20.0 | 21.0 | 22.0 | μM |
| hygromycin B | unknown | 0 | 10.0 | 15.0 | 20.0 | μM |
| nocodazole | TUB1 | 0 | 11.0 | 12.0 | 13.0 | μM |
| cytochalasin A | ACT1 | 0 | 40.0 | 50.0 | 60.0 | μM |
| pentamidine | unknown | 0 | 75.0 | 115.0 | 130.0 | μM |
| nigericin | unknown | 0 | 20.0 | 25.0 | 30.0 | nM |
| neomycin sulfate | unknown | 0 | 175.0 | 200.0 | 225.0 | μM |
| sodium butyrate | HDACs | 0 | 40.0 | 60.0 | 80.0 | mM |
| 3470-5652 | unknown | 0 | 9.0 | 11.0 | 13.0 | μM |
| 1273-0059 | unknown | 0 | 1.4 | 1.7 | 2.0 | μM |
| 3770-0098 | unknown | 0 | 150.0 | 200.0 | 250.0 | μM |

### Predicting synergy

“Chemical space — a term used to encompass all possible small (>500 atoms) organic molecules, including those in biological systems — is vast”^41^. Furthermore, the current purchasable, and readily screenable compounds comprise approximately 8 million unique compounds^42^. Screening each of these molecules as single agents is quite daunting, while screening all pairwise combinations is impossible. The number of potential combinations is (N^2^ − N)/2, with the number of starting compounds being (N). To expedite the identification of synergistic combinations, we test if we could uncover synergistic combinations from combination chemogenomic data.

We reasoned that if a drug induces a fitness defect in a particular gene-deletion mutant, but does not directly inhibit that gene product, then this drug might be synergistic when combined with a second compound that does inhibit that gene product. To survey the possible drugs and mutants that satisfy these criteria, we first used our database of several thousand chemogenomic assays^14^ to define when a heterozygous deletion mutant of a known drug-target is sensitive. We then sought to uncover synergistic interactions so the drug can be paired with a second drug that inhibits the known drug-target (Figure 3).

**Figure 3.**
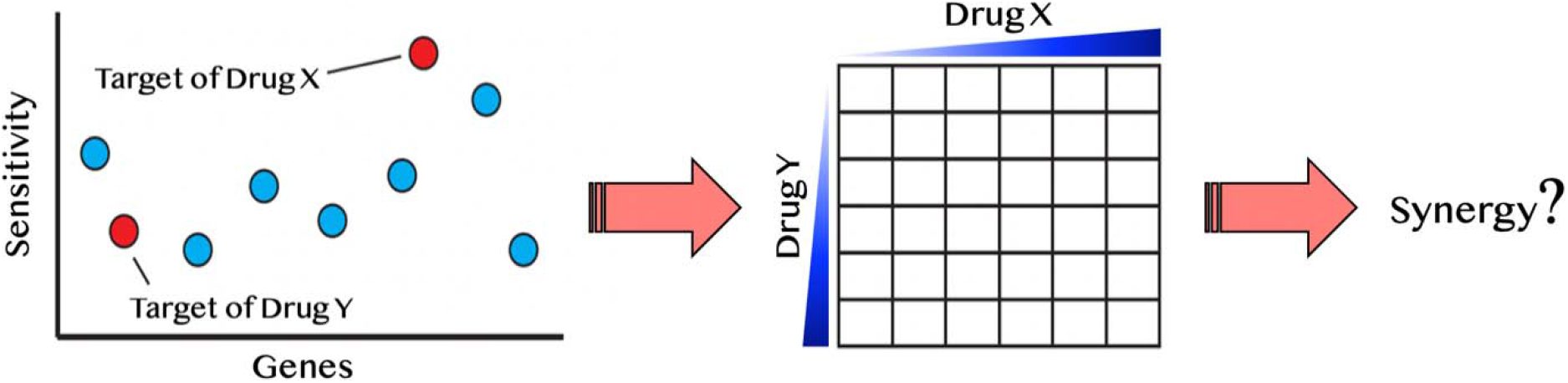
Diagram of synergy prediction method. The HIP–HOP profile for drug X, in which red circles are essential genes and blue are nonessential genes. Genes are listed on the x-axis and sensitivity to drug X is on the y-axis. The most sensitive strain in this example is the target of drug X. However, another essential gene also displays sensitivity to drug X. This gene is the known drug-target of drug Y. Using the synergy prediction method, drug X and drug Y would be predicted to be synergistic. By testing them in a drug dosage matrix, one can determine if this assumption is true.

This principle is illustrated with miconazole, an antifungal that targets the enzyme, Erg11p. Because the hmg1Δ deletion strain is also sensitive to miconazole, we hypothesized that the combination of miconazole and an HMG1 inhibitor, (e.g., cerivastatin) would be synergistic. To test this hypothesis, we first examined the dataset in reference^43^ and all single-agent screens performed in our lab to ask if any of the drug targets in Table 1 exhibited a fitness defect. From this survey, 25 predicted synergistic combinations were selected and empirically tested to determine if their synergy rate was greater than the background synergy rate of 17%. We found 40% of the tested pairs were synergistic (ε < −0.20): a 2.5-fold significant enrichment over random pairs (p-value < 0.01). This approach is conceptually distinct from another synergy prediction method introduced by Jansen et al.^44^ which computed a correlation between HIP–HOP profiles. Our approach in contrast, is empirical, asking if a drug, selected based on its drug-gene interactions in HIP–HOP, can induce synergy. This approach is well-suited for profiles with low similarity. By way of example, miconazole and fenpropimorph, is a synergistic combination that we identified despite a low profile similarity^44^.

### HIP–HOP Combination Profiles

We next used a variation of the HIP–HOP assay, testing compound combinations genome-wide. All 14 confirmed synergistic combinations and 12 non-synergistic combinations, and each single agent was used for chemogenomic screening. We then used the approach described by Lee et al.^14^ to identify any significantly sensitive strain in all screens. To further scrutinize the drug combinations, we defined the epsilon, ε, as a fraction of uniquely sensitive genes, in both synergistic and non-synergistic pairs. We defined uniquely sensitive genes as those that are significant only when both compounds are tested together at a fitness defect score of 2.0 or greater. Two examples are shown in Figure 4, and the entire dataset of single agent and combination HIPHOP profiles is in supplementary information (synergy files).

**Figure 4.**
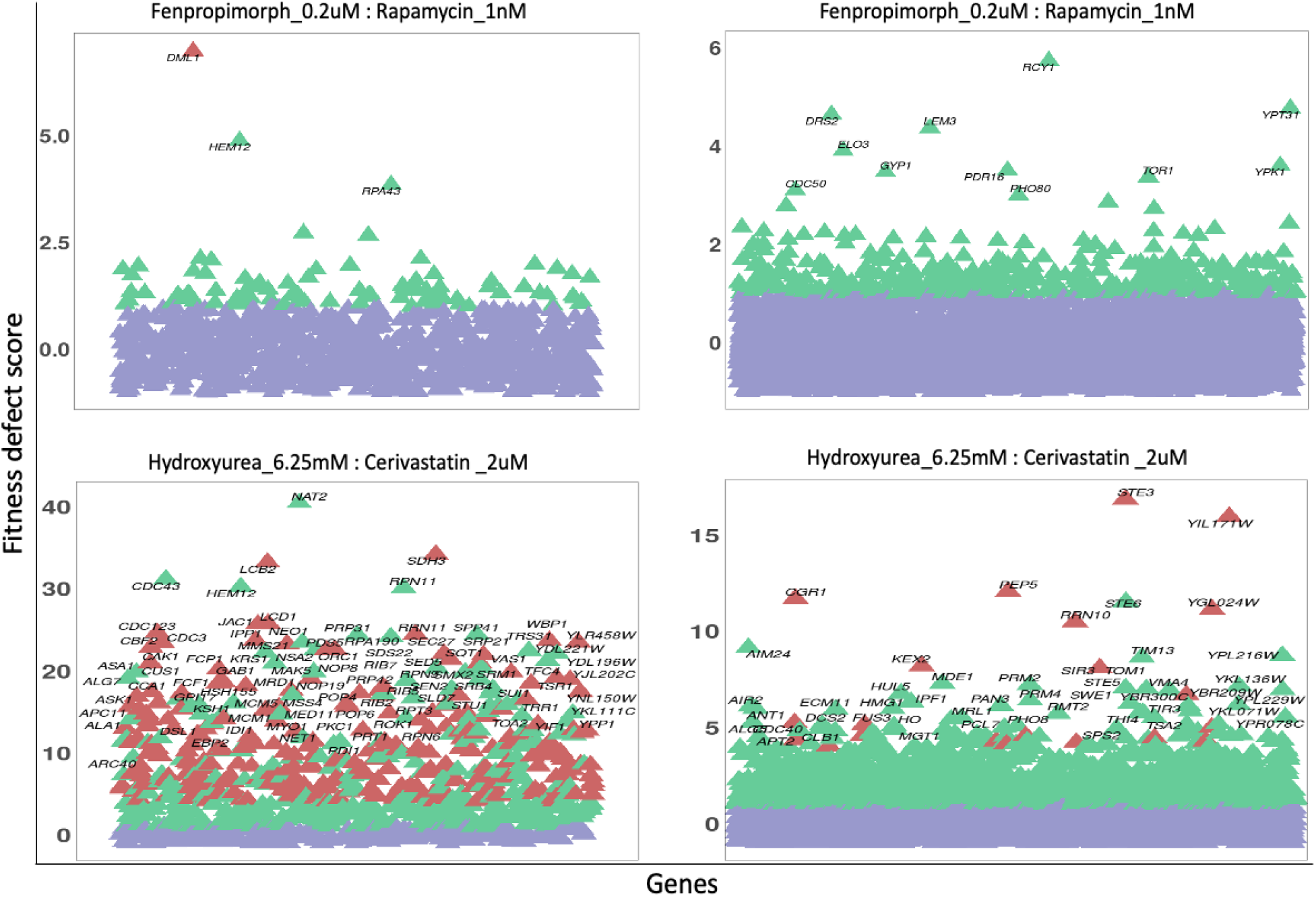
HIPHOP screening of select drug combinations. For each combination screen, a drug combination that inhibited WT yeast growth by 20% was selected and screened alongside each single agent. To identify combination-specific strains, we required that the fitness defect in the combination be 4.0 or greater and that in each single agent, the fitness defect for that strain did not exceed 2.0. For clarity, the heterozygous essential (HIP- left scatterplots) and homozygous non-essential (HOP- scatterplots) data are plotted separately. Significantly sensitive strains are highlighted in green, and combination-specific strains are depicted in red, with the fitness defect scores shown on the y-axes. In the case of the fenpropimorph x rapamycin combination (top plots), only a single strain- the essential gene DML1. This gene product has been implicated in diverse aspects of mitochondrial function. For the hydroxyurea x cerivastatin combination (bottom plots), a larger number of combination-specific strains are apparent. Among these are essential genes involved in sphingolipid biosynthesis (LCB2), mitochondrial metabolism (SDH3, JAC1) as well as cell cycle checkpoints, and protein degradation at the metaphase anaphase transition (LCD1, CDC23, and CBF2). Non-essential strains specific to this combination include those involved in response to diverse stresses (STE3, CGR1) and targeted protein degradation (PEP5, KEX2).

Although, the primary goal of these genome-wide combinations is to serve as a resource for focused tests of individual combination-specific genes, several high-level observations are noteworthy; i) combinations vary greatly in the number of specifically sensitive genes, ii) in some cases the combination-specific strains appear to be subject to potentiation by one of the two agents (i.e., these strains can be detected at higher doses of the single agents^45^, and iii) the combination-specific genes identified are consistent with known mechanisms of actions of one or both of the drugs used in the combination.

We examined each combination screen, both synergistic and non-synergistic, and examined the biochemical pathways enriched in each pathway. Examining the Gene Ontology (GO) enrichments via the synergy score (Figure 5), we found that each combination provides a unique signature. For instance, the miconazole-cerivastatin combination screen was enriched for gene deletion strains involved in cell wall, cytokinesis, vesicle-related processes, and sterol biosynthesis. In contrast, the miconazole-hydroxyurea screen is enriched for vesicle-related processes and cytokinesis but not for cell wall-related or sterol biosynthesis processes. These combination-specific GO enrichments can identify which cellular processes are providing resistance to the combination and could help to understand the mechanism of synergy on a combination-specific basis.

**Figure 5.**
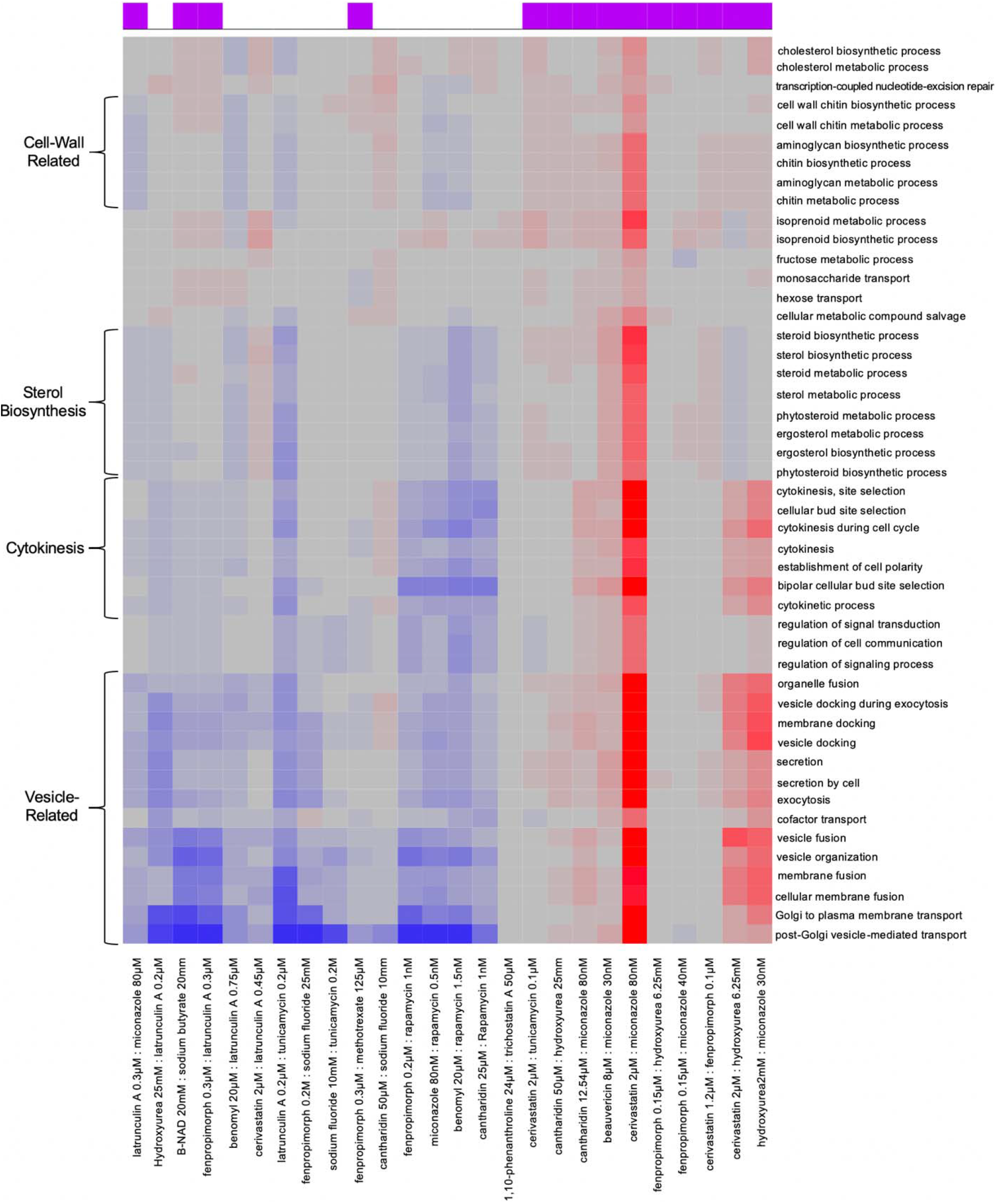
Combination-specific Gene-Ontology (GO) enrichment HIP–HOP data. Examination of Gene-Ontology enrichment using ε score. In the clustered heat map generated from the ε scores, one the x-axis we have each drug combination listed on the bottom; at the top on the x-axis is a purple box which denotes if a combination is synergistic or not. GO terms are denoted on the y-axis. Red shows a particular GO term is highly enriched in the combination, grey denotes no enrichment and blue shows significant enrichment among genes with low ε scores.

## Discussion

In this study, we used genome-wide chemogenomic profiling to select drug combinations for synergy testing then confirmed our predictions using combined chemogenomic assays. An interesting observation from drug combination data is that the three inhibitors that affect the ergosterol pathway were highly synergistic when applied in combination, suggesting that compounds that inhibit different points within a pathway are more likely to be synergistic, consistent with Zimmermann et al.^46^ and the observations by Cokol (2011) that found similar compounds to be “promiscuously” synergistic^12^. We further demonstrated that, among the synergistic combinations, 78% of the combinations tested (Table 1) were the result of combining an ergosterol inhibitor with a second agent. This indicates that the ergosterol inhibitors are highly synergistic with other agents, which is likely due to their effect on the yeast cell membrane, thereby allowing compounds more effective entry^40^.

Using a modified genome-wide assay we demonstrated that synergistic combinations result in uniquely sensitive strains that are specific to the combinations and are not observed in either of the single agents. Because we used stringent cut-offs, the difference we found between synergistic and non-synergistic combinations likely represents a minimum level of enrichment. We also found that each combination has its own pathway and GO enrichments (Figure 5).

During the course of this work, confirmed that these drug-drug interactions (derived from drug combination treatment) can be recapitulated using drug-gene interactions by directly assaying loss-of-function (heterozygous deletion) mutants for a drug’s known target with a drug that inhibits a synergistic target. We further found that drug-gene interactions derived from synergistic drug-drug interactions were enriched for negative interactions. To extrapolate these observations, we analyzed our single-agent chemogenomic screening data to predict combinations that might exhibit synergy. Given that we observed the baseline level of synergy between 10-17%, between 83-90% of any random combination should not be synergistic. Our approach reduces the screening required by at least 2.5-fold. This experimental approach involves; 1) using chemogenomic data to identify drugs able to make known drug-targets haploinsufficient, 2) pairing the strain with the expected drug, and 3) screening a dose matrix for synergy. In a pilot of 25 combinations, we identified 10 synergetic combinations This method is easily adaptable to include new drug-targets, as we limited our search to 11 well-characterized drug-targets in Table 1 and only examined one dataset^43^.

Synergistic effects (between either genes or drugs) have received renewed attention, especially in light of increasingly sophisticated computational approaches and the precise genome engineering possible with CRISPR-based technology. For example, Cokol et al. developed a computational framework called MAGENTA to investigate the impact of microenvironment on antibiotic combinations, stating that it enables tailoring antibiotic therapies based on the pathogen microenvironment. For MAGENTA to predict synergistic or antagonistic interactions on various microenvironments, it leverages chemogenomic profiles of both single drugs and metabolic perturbation. They reported several synergistic combinations against E. coli and A. baumannii, and predicted bactericidal drug-combinations' effectiveness when grown in glycerol media and classified genes in glycolysis and glyoxylate pathway as top predictors of synergy and antagonism, respectively^47^.

In 2016, Wong et al. leveraged combinatorial genetics en masse (CombiGEM) to systematically study gene and drug combinations modulating biological phenotypes^48^. CombiGEM allows for the rapid construction of barcoded, combinatorial genetic libraries that can be quantified with high-throughput sequencing. They applied CombiGEM-CRISPR to generate a library of 23,409 barcoded dual guide-RNA (gRNA) combinations, performing a high-throughput pooled screen to find gene pairs that combine to inhibit ovarian cancer cell growth. In the same study, small-molecule drug pairs were tested against the pairwise synthetic lethal hits, revealing that they exert synergistic antiproliferative effects against ovarian cancer cells^48^.

Combining chemogenomics and genetic interactions, Weinstein et al. studied antifungal combinations applied to two yeast species, C. albicans and S. cerevisiae. This study showed that, both synergistic and antagonistic combinations increase the cell-type selectivity of growth-inhibiting drugs. The authors speculate that drug interactions might shift selectivity in comparison to single-drug effects in mixed microbial communities. Indeed, few drugs or drug combinations should be expected to encounter the idealized conditions in laboratory experiments- the variations observed by Weinstein^49^ can change the selectivity of compounds, i.e., inverting, diminishing, or enhancing therapeutic windows.

In a recent CRISPR/Cas9 screen Huang et al. sought to identify genes whose depletion causes synthetic lethality with the broad-acting but not particularly potent Aurora kinase inhibitor VX-680^50^. They reported that HCT116 cells showed hypersensitivity to VX-680 when Haspin— a serine/threonine-protein kinase encoded by GSG2 gene — was either depleted by CRISPR knockout or with Haspin inhibitors, confirming the synergistic effect between VX-680 and Haspin depletion or inhibition^50^. Recently, Zhou et al. reported a CRISPR-based, multi-gene, knockout screening system for assembly of barcoded, high-order combinatorial guide RNA libraries, en masse. Although combination therapies promise to improve treatment efficiency of various diseases, only a few effective drug combinations — especially those employing three or more drugs (see table S1 in reference^51^) — have been introduced so far. Zhou et al. used this approach to systematically identify both pairwise and three-agent synergistic therapeutic target combinations. The study claimed to uncover double- and triple-combinations that suppressed cancer cell growth and afforded protection against Parkinson’s disease-associated toxicity^51^.

## Conclusion

In this work, we introduce a strategy to use comprehensive genome-wide screens to first predict compounds that might be synergistic and then test novel combinations empirically. This approach should be extensible to other models and allow for a rational approach to selecting effective drug combinations.

## Methods

### Pair-wise screening of drug combinations

To identify synergistic combinations, one should determine the effect of both individual agents and drug combinations on yeast growth rate. To accomplish this, we screened the drugs in a checkerboard matrix, in which, along each axis, one of the drugs is added at progressively higher doses. Drug concentrations were selected based on inhibitory concentration (IC), which was determined prior by prescreening the drugs’ effects on the wild-type cell growth; concentrations are such that there is an IC0 (no drug), IC2, IC5, IC10, IC20, and IC50 for each drug in the matrix. Hence, each drug pair was screened in a 6-by-6 dose-response curve at the IC values listed above. All possible pairs of the 11 drugs were screened, resulting in 55 combinations, including self-by-self drug combination. The yeast strain BY4743 (S288C) was used to screen these combinations and was grown in a 96-well microtiter plate at 30°C for 24 hours in a TECAN optical density reader. The optical density (OD_600_) was measured at 15-minute intervals. A diagram depicts how the combination screening was performed (Figure 1).

### Determination of synergistic combinations based on growth curve analysis

Following growth, data for all growth curves were extracted using AUDIT software^15^ as described. First, the curves were smoothed, and the area under the curve was calculated. The area under the curve was then compared to the area of no drug control (AREA_Drug_/AREA_No Drug_) to create an inhibition ratio. We then used the Bliss multiplicative model^3^ to calculate epsilon for each dose matrix, ε = Drug AB_Ratio_ – (Drug A_Ratio_ × Drug B_Ratio_). Specifically, we considered “drug epsilon” to be the different between the actually combined growth and the ‘expected’ which the multiplication of the two single agents. For example, if Drug A grew at 90% compared to no drug and Drug B grew at 80% compared to no drug, the expected defect would be 90% × 80% (e.g., 72%). If the actual combination grew at 50% compared to no drug then elision would be 50% - 72% = −23% When epsilon is zero, then no interaction is observed; when epsilon is negative, there is a synergy, and positive epsilon denotes antagonism. An antagonistic interaction indicates that one of the drugs buffers the effect of the second agent— Fig. 1 illustrates how growth data was transformed into a quantitative trait to determine epsilon.

### Drug-gene interaction screening using isogenic cultures

To determine whether a deletion mutant was hypersensitive to the drug, we had to know growth rate of both the mutant and wild-type strains with and without drug. We used heterozygous deletion mutants of the known drug-targets (listed in Table 1). The yeast strains BY4743 and corresponding heterozygous mutants were grown, as isogenic cultures, in 96-well microtiter plates, at 30°C for 24 hours in a TECAN optical density reader. The optical density (OD_600_) was measured at 15-minute intervals. Here, the growth metric average generation time (aka AvgG) was used to assess the fitness of wild-type and mutant strains with and without drug; this metric is comprehensively described in the protocol written by our lab^52^. We normalized each strain’s fitness to the wild-type and subtracted any single mutant fitness that was contributed by any particular mutant, i.e., we normalized the various heterozygous mutants’ growth to wildtype to take into account any fitness defect that was caused by haploinsufficiency.

### Predicting synergistic combinations via chemogenomic interactions

Following up on how drug-drug interactions predict drug-gene interactions. To predict synergy using chemogenomic data, we examined 18 datasets (see Table 2) and assessed if the known drug-targets listed in Table 1 were sensitive in any of the treatments based on the log_2_ ratio of control over treatment. We then identified combinations available in our laboratory for testing; 25 combinations were selected based on the availability of compounds present in the Giaever/Nislow laboratory as well as Boone lab, at the University of Toronto. To determine if this method can successfully predict synergistic combinations, the chances of observing synergy between randomly paired compounds need to be known.

### Determination of background synergy rate and experimental validation of predicted combinations

To define enrichment for synergistic combinations, the chances of observing synergy between randomly paired compound combinations must be known. To address this, 105 combinations in a 4-by-4 dosage matrix were screened. We used a smaller matrix in this experiment to maximize the number of combinations that could be screened in a short time. As a result, 6 combinations can be screened per 96-well plate per TECAN plate reader, instead of 2 combinations when screened in a 6-by-6 matrix. The drug concentrations were such that there was an IC0 (no drug), IC10, IC20, and IC50 for each drug in the matrix (Table 2). The yeast strain, BY4743, was used to screen these combinations and was grown in a 96-well microtiter plate at 30°C for 24 hours in a TECAN optical density reader. The optical density (OD_600_) was measured at 15-minute intervals.

### Pooled competitive growth assays

Two deletion pools, a homozygous deletion pool of 5054 strains representing non-essential genes and a heterozygous pool of 1194 strains representing genes essential for viability, were thawed and diluted in YPD to an OD_600_ of 0.0625; 700 μL cultures were then grown at 30°C with a chemical inhibitor(s) applied at a dose that produced 10-20% growth inhibition of wild-type. An automated liquid handler robot was used to maintain logarithmic growth of pools by collecting 0.7 OD_600_s of heterozygous pool following 20 generations of growth, and 1.4 OD_600_s of homozygous pool following 5 generations of growth, for further processing as described below.

### Assessing fitness of barcoded yeast strains by barcode microarray

Except where indicated, pooled assays were performed as described in the protocol by Pierce et al^53^. Genomic DNA was isolated from cells and barcodes, amplified, and hybridized to barcode microarrays, where each barcode deletion mutant is represented by ten hybridization signals (the uptag and downtag for each strain are each represented on the array five times)^53^. Array measurements were quantile normalized such that all tags hybridized with the sample pool had similar distributions. Following normalization, we applied a correction factor to the array data to correct for feature saturation^53^ and determined the fitness of each barcoded deletion strain using the average of both barcodes. A Z-Score was calculated based on the average barcodes signal intensity against a control probe sets distribution. Positive fitness defect scores signify a decrease in strain abundance during drug treatment.

### Haploinsufficiency profiling (HIP) and Homozygous profiling (HOP) of synergistic combinations

A key parameter in performing genome-wide screens in yeast is to determine an appropriate screening dose. This value has been empirically determined to be 10-30% of inhibition of wild type growth^14^. In practice, when performing synergy screens with two agents, one must eliminate any effects due to the action of a single agent alone. Therefore, we screened each single agent at its IC20, as well as at the dose that was used to generate a combined IC20. We therefore needed data from both the combination and individual agents. For each combination genome-wide assays 5 screens were performed. Specifically, Agent A at its IC10-30, Agent A the dose used when combined (usually an IC2), along with Agent B at its IC10-30, Agent B at the dose used when combined (usually an IC2). Accordingly, each genome-wide synergy assay comprises five separate screens; 1) the combined screen (A+B), 2,3) each single agent at an IC10-30 (A and B, 4,5) each single agent at the dosed used in A+B.

### Analysis of combination profiles

#### Defining the sensitivity score

For each drug treatment, a Z-Score based on the averaged Up and DOWN barcode signal intensity was calculated. Using a Z-Score of greater than 4, we defined a list of sensitive strains in each treatment. Filtering out strains that arose from the single drug treatments, we were able to identify unique, combination-specific sensitive strains. To aid in the analysis of drug combinations, we defined factor ε that is the sensitivity of genes in the combination minus the sum of sensitivity in the single agents — ε = Z-Score_AB_ – (Z-Score_A_ + Z-Score_B_).

#### Clustering of combination HIPHOP profiles sensitivity scores

We took raw intensity values from the barcode microarrays and normalized the logged raw intensities using a method called Supervised Normalization of Microarrays (SNM). This method^54^ was supplied with batch definitions — each batch as the arrays that have the same chip date. We also supplied this method with array descriptors corresponding to the chemical treatments (i.e., which compounds were used); this is meant to preserve biologically relevant signal. After SNM, we selected either the uptag or downtag for each strain based on the lowest variation coefficient to avoid noisy tags. The logged intensities were then used to compute robust Z-Scores. We used the median and median absolute deviation to calculate the Z-Score, and then clustered strains and chemical treatments separately. The similarity between strains/chemical treatments was based on the Pearson correlation of Z-Scores.

#### Examination of gene-ontology terms

Gene-Ontology has the aim to standardize terms for describing gene products. This vocabulary defines a set of cell terms for which a gene can be annotated to. These annotations cover a vast range from location within the cell to specific cellular functions such as nucleotide excision repair. In this study, we used Gene-Ontology terms with more than 5 genes and less than 200 genes. To determine enrichment, we used the sensitivity score. Following ranking each gene sensitive in a specific combination, we used Gene Set Enrichment Analysis (GSEA)^55^ to determine enrichment in each category.

## Acknowledgements

We thank Iain Wallace for advice and reading early drafts of this work and Marjan Barazandeh for experimental assistance.

